# Ontogeny of tactile, vocal and kinship dynamics in rat pup huddling

**DOI:** 10.1101/2025.04.08.647436

**Authors:** Florbela da Rocha-Almeida, Hugh Takemoto, Ann M. Clemens

**Author notes:** Corresponding author: Ann Clemens.

## Abstract

Huddling, a tactile, thermoregulatory and filial social interaction, is a predominant and conserved social interaction mammalian and bird species. The act of huddling is particularly important in early life, when thermoregulation, social touch and bonding are influential for survival and healthy brain and behavioural development. We ask how tactile, vocal and kinship dynamics of rat pup interactions develop. We designed a huddling apparatus where we record and synchronise huddle formation with ultrasonic vocalisation analysis. With development, we see that groups (P6–8 vs older pups) stay longer in triad aggregon (pup huddle) configurations in the huddle trial period. Older pups (P18–20) switch huddle states more often. The spectral characteristics of rat pup vocalisation during huddling task change in development, with a higher peak frequency in P18–20 pups. In all age-groups we observe vocal quieting as aggregons form. We hypothesized that kinship should be a strong determinant of huddling interactions but our findings reveal otherwise. When comparing kin vs non-kin groups we found no differences in aggregon durations or switches. In the youngest age group (P6–8) the amount of USVs were reduced in kin vs non-kin groups, though not in older age groups. To address the role of tactile contact in quieting we integrated touch dividers in the huddle arena. Without touch, we saw that vocalisations significantly increased in P6–8 and P11**–**14, but not in P18–20 kin sibling groups. We suggest that rat pups have a strong internal drive towards huddling behaviour regardless of whether huddle partners are related by kinship. USV analysis suggests that huddling has a calming effect, where related sibling young pups show less distress overall; absence of touch is associated with increased distress in P6–8 and P11**–**14 kin huddle groups. Thus, huddling is a natural social behaviour that is intrinsically rewarding and shared between both related siblings and unrelated conspecifics; it has calming effects - as indicated by USVs - that depend on kinship and tactile contact during the earliest stages of development.

## Introduction

Social contact and warmth, through thermoregulation or emotional support, are important factors in mammalian development which influence later-life mental health and survival [1]. The importance of conspecific contact in development has been emphasised by severe consequences that lack of social touch has on well-being and health [2–4].

Huddling behaviour features predominantly in mammalian and bird species, where huddling can reduce metabolic costs for individuals and enhance survival of the group. Emperor penguins, for example, utilise huddling to survive prolong fasts and ensure the survival of offspring [5,6]. Maternal touch has been identified as an important early life stimulus, however sibling touch can be as prevalent, if not more prevalent, in early life experience. Rabbit pups which huddle with siblings conserve more energy than individually reared animals, leading to more conserved energy for nursing and milk consumption [5,7].

In rodents, huddle preferences show species diversity, where Prairie voles, but not mice show preference for huddling with social partners and familiar individuals [8,9]. Rat pup huddling functions in early development for thermoregulation and later becomes an olfactory-based filial activity, indicating a role in social bonding beyond metabolic conservation [10,11]. Previous work indicates that rat pups may show kin preference in huddling [12]; studies in Prairie voles indicate partner and familiarity preference in huddling [8,9].

Rodents elicit ultrasonic vocalisations (USVs) which express the internal state of the animal [13]. Rat pups elicit distress calls with separation, temperature discomfort or anxiety [14–16]. In older pups, USVs may indicate positive emotional states such as in play behaviour [17]. Calling interactions between the mother and pup have been well characterised, and the neural circuits described [18], however vocalisation between peer and sibling animals in development, specifically in the context of pup huddling, is less understood.

To address the hypothesis that huddling and vocal dynamics of rat pups are regulated by development, sex and kinship, we designed a setup for recording and quantitatively analysing rat pup tactile and vocal interactions in natural huddling behaviour.

## Materials and Methods

### Animals

Long Evans outbred rats were bred and maintained on a 12-hour/12-hour reverse light-dark cycle and provided *ad libitum* access to food and water. All animal procedures adhered to the regulations outlined by the United Kingdom Home Office. For each experiment, three animals were randomly selected per group (male kin, male non-kin, female kin, female non-kin), and animals were only used once for behavioural experiments to eliminate the possibility of adaptation. The total number of groups of animals was 30 groups for postnatal days P6 – P8; 29 groups for P11 – P14; and 31 groups for P18 – P20. Two experimental sessions were excluded from the vocalization analysis due to the lack of TTL (transistor-transistor logic) signals required for alignment with video data. Outbreeding of rats was maintained locally by tracking genealogies and by importing novel animals from the supplier to refresh stock (Charles River, Italy).

### Huddling behaviour setup

An open arena made of flexible plastic, measuring 22 cm in diameter and 10 cm in height, was utilized for P6 – 8 and P11 – 14 sessions, while for P18–20, the arena dimensions were increased to 33 cm in diameter and 31 cm in height. Experimental conditions were maintained in a dimly lit room with controlled temperature (approximately 18–21°C). Video recordings and ultrasonic vocalizations were captured using IC Capture software (Imaging Source monochrome camera) and Avisoft ultrasonic vocalisation recording (USV) software, respectively.

### Video Analysis

Video analysis was conducted blinded to group conditions using Elan software (version 6.3). Vocalizations were analyzed using DeepSqueak and alignment of vocalisations and behavioural scoring was done with Matlab. Aggregon number shows the quantity of animals huddling. Aggregon 0 represents complete separation of all three animals, while aggregon 2 indicates two animals huddling together, and aggregon 3 signifies three animals huddling together.

### Aggregon switches, Aggregons and Calls Per minute

For each minute of the 20 min session, the number of aggregon switches were counted. The total number of switches were then summed across the 20-minute session to extract the total number of aggregon switches. To quantify aggregon score per minute, 1 was assigned to aggregon 0, 2 to aggregon 2 and 3 to aggregon 3. Aggregon score per minute was assigned by scoring the video frame at 1-minute intervals. The score across groups was then taken as the average for all groups per minute. To quantify calls per minute, the number of calls were summed in 1-minute intervals. The mean for each minute was then taken across all huddle groups.

### Statistical Analysis and model

Groups were compared without assumptions of normal distribution, thus non-parametric testing was applied (Mann-Whitney, Kruskal-Wallis tests). Data are presented as mean ± SEM unless stated otherwise. To create the probability matrix used for the Markov chain model, a probability was calculated for each state possibility from the 1200 states (one per second) which were scored for each group. The mean of the probabilities across groups was then calculated and used in the model. The model was then run 25–30 times to replicate group variability across 1200 steps (the length of the experimental recording). The initial state was set to 1 (indicating aggregon 0) as was the convention of the scoring scheme to create the aggregon scores of the data.

## Results

### Experimental setup for examination of huddling behaviour development in rat pups

To address the organisation, development, vocal and kinship interactions which occur throughout development, we designed a huddling apparatus in which we monitored interaction in triads of rats (male kin, male non-kin, female kin, female non-kin) at three pre-weaning developmental stages (P6–8, 11–14, 18–20). The size of the arena was modified to accommodate the size and mobility of the rat pups. The arena was illuminated with infrared illumination and filmed with a monochrome camera which was synchronised to the ultrasonic vocalisation recording (**Figure 1A**). In each developmental stage, rat pup groups were placed in the arena in a separated state in a triangular configuration (aggregon 0) at the start of the trial and pups were allowed to behave freely. The trial period was limited to 20 minutes. Kin groups consisted of three male or female siblings or non-kin groups (one from each litter of a timed-mated female mother). Rat pups were used for only one trial and were not re-used. We scored contact behaviour as aggregons where an aggregon of 0 consisted of no pups huddling, aggregon of 2 consisted of 2 pups in contact and aggregon of 3, where all pups were in a huddle (triad) (**Figure 1B**). In the example trial we found that rat pups alternated between aggregon 0 and aggregon 2 before forming a triad (aggregon 3) and largely remaining in an aggregon of three with few alternations back to aggregon 2. Aggregons for the entire session were scored and aligned to vocalisations using synchronisation methods (**Figure 1C**). Vocalisations occurred more frequently at the start of the twenty-minute trial which then reduced over the trial time (**Figure 1D**).

**Figure 1.**
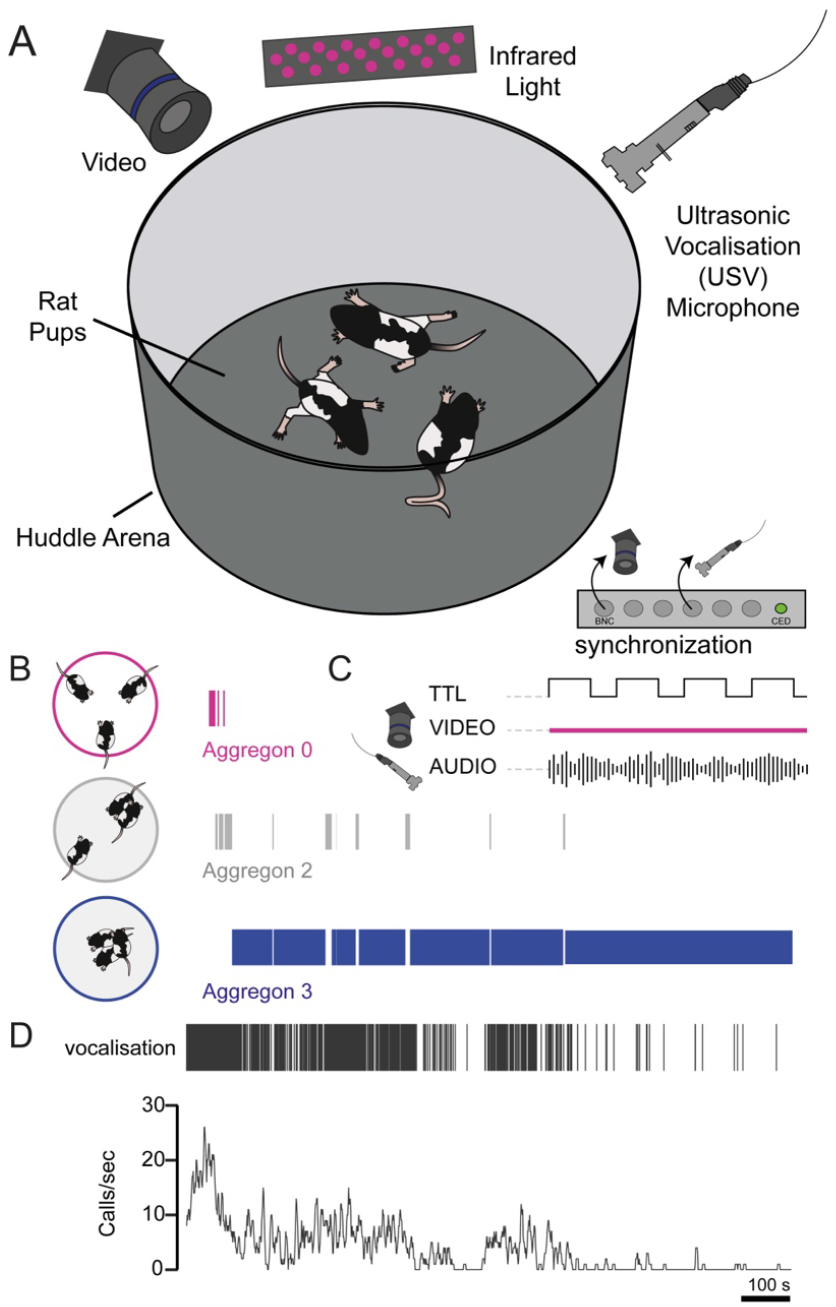
Experimental setup for examination the development of huddling behaviour in rat pups. A) The huddling setup consisted of a circular arena which was placed upon a washable base inside a behavioural cabinet which was sound-insulated and dark. Video recording was performed with infrared illumination and recording of ultrasonic vocalisations with one microphone. Temperature was monitored and maintained consistently at 18–21°C. B) Three rats were placed in a triangular configuration at the start of the experiment. A typical progression of aggregons is displayed, where separated rats (aggregon 0) transitioned to an aggregon of 2 (2 out of 3 rats touching) and then progressed further to all rats huddling (aggregon 3). Aggregon states fluctuated throughout the experimental time (20 minutes total). C) Video recording was synchronised with ultrasonic vocalizations (USV) with a Cambridge Electronic Design (CED) board which triggered camera frames and elicited an electronic (TTL) time stamp data which was encoded in the USV audio file via Spike2 software (CED). D) Representative USV events which were detected using DeepSqueak, deep-learning based analysis package for USV spectrograms. Vocalisation frequency was initially high in the huddle trial and reduced in frequency as the huddle trial progressed.

### Aggregon configurations increase in complexity as rat pups develop

We next addressed the progression of aggregons across twenty-minute trials with developmental days. When placed in the triangular start formation, pups transitioned between aggregons 0, 2 and 3 (**Figure 2A**). Huddle progression was visualised across groups (top P6– 8, middle P11–14, bottom P18–20) for the twenty-minute trial (**Figure 2B**). All aggregon configurations were found to vary with development. Triad (aggregon 3) duration was the least occupied state at P6–8 and the most occupied state at developmental stages P11–14 and P18–20. (**Figure 2C**). Thus, as pups develop, aggregon complexity increases.

**Figure 2.**
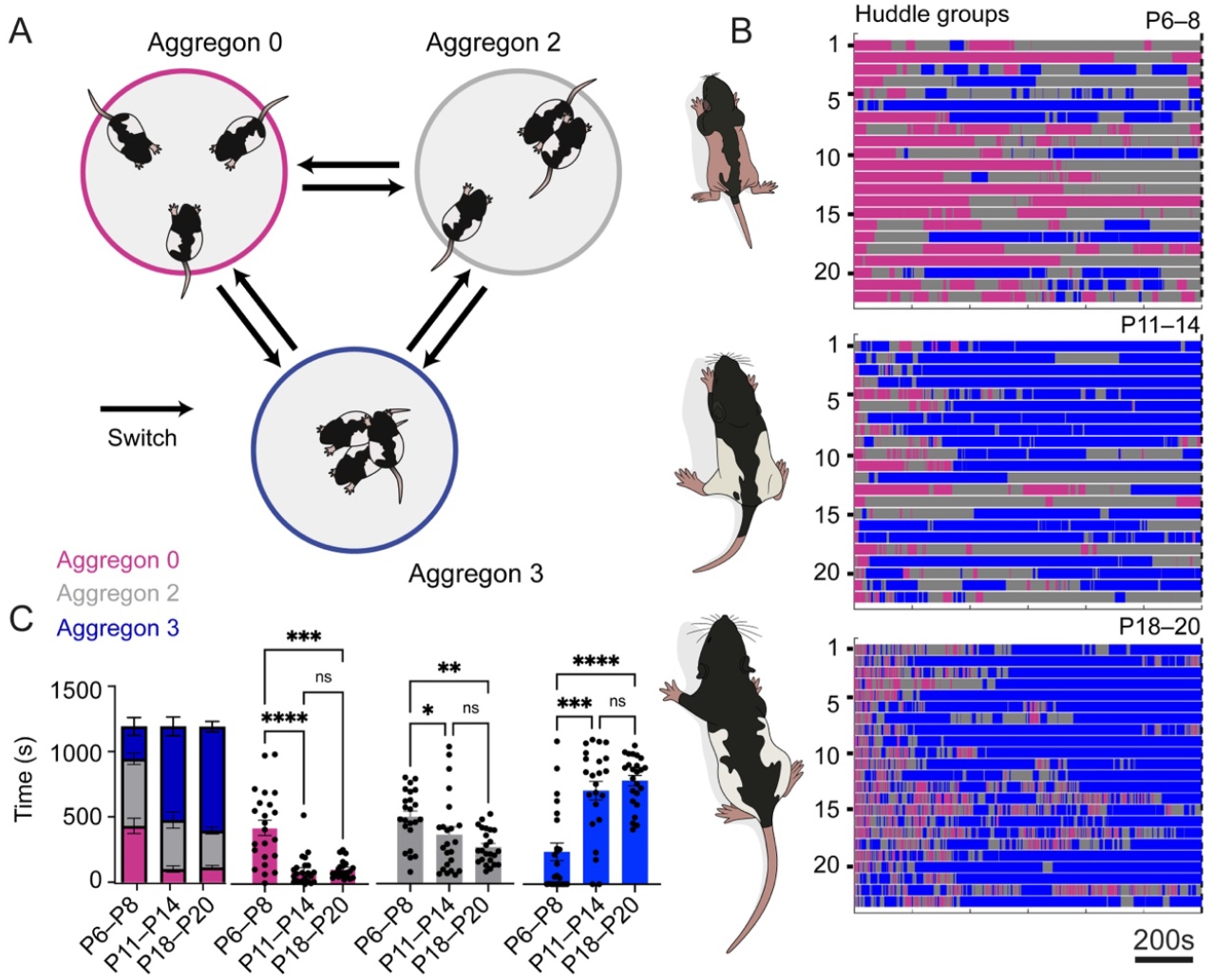
Aggregon configurations increase in complexity as rat pups develop. A) Depiction of aggregon 0, 2 and 3. Transitions can occur between all aggregon states. B) Raster of aggregon groups. Aggregon 0 is depicted in magenta, aggregon 2 in gray and aggregon 3 in blue. Total time displayed is 20 minutes, scale bar 200s. C) Quantification of total time spent in aggregon state throughout development. Individual data points represent groups of 3 rat pups. Magenta: Aggregon 0 durations vary with development (Kruskal-Wallis test, p < 0.0001). Time spent in aggregon 0 is highest P6–8 (P6–8 vs P11–14, p<0.0001; P6– 8 vs P18–20, p=0.0004), without a significant difference between P11–14 and P18–20 (p= 0.9365). Gray: Aggregon 2 durations vary with development (Kruskal-Wallis test, p = 0.0032). Time spent in aggregon 2 is highest P6–8 and reduces with development (P6–8 vs P11–14, p=0.0285; P6–8 vs P18–20, p=0.0018), without a significant difference between P11–14 and P18–20 (p > 0.999). Blue: Aggregon 3 durations vary with development (Kruskal-Wallis test, p<0.0001). Time spent in aggregon 3 is highest P6–8 and reduces with development (P6–8 vs P11–14, p=0.0003; P6–8 vs P18–20, p<0.0001), without a significant difference between P11–14 and P18–20 (p>0.999). Kruskal Wallis followed by Dunn’s multiple comparisons test (N=23 P6–8, N=23 P11–14, N=24 P18–20). Left axis corresponds to all graphs. Bars indicate mean ± SEM.

### Huddle transitions are regulated by developmental age

We observed that rat pup groups occupied three main states (aggregon 0, 2, 3) and alternated between these three states in all developmental stages. To examine whether the number of transitions between aggregon states was developmentally regulated, we summed the number of aggregon switches throughout the twenty-minute trial. We found that the number of switches varied across age groups with significant increases between P6–8 and P18–20, and P11–14 vs P18–20 (**Figure 3A–B**).

**Figure 3.**
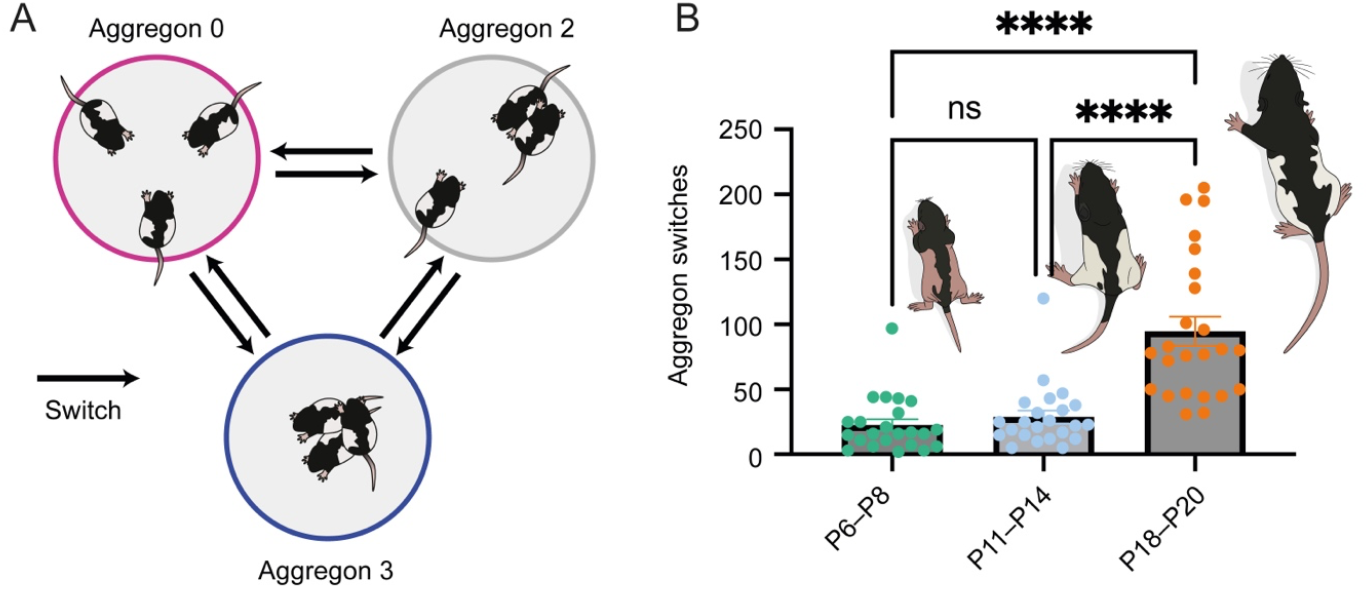
Aggregon switches increase with post-natal development. A) The number of aggregon switches across the 20 minute session were summed. For each minute of the video, the number of aggregon switches were counted. A switch was defined as a transition between aggregon 0, aggregon 2 or aggregon 3, in any direction. 3) The number of total aggregon switches across all groups (P6–8, 22.61 ± 4.40; P11–14, 28.83 ± 5.017; P18–20, 94.88 ± 11.16 switches Mean ± S.E.M.) increased with developmental age. Number of switches varied with developmental age (N=3 age groups, p<0.0001, Kruskal-Wallis test). Switches for group comparisons was not significantly different between P6–8 and P11–14 (p>0.999, N=23 P6–8, N=23, P11–14) and was significantly different in P6–8 vs P18–20 (p<0.0001, N=23 P6–8, N=24 P18–20) and P11–14 vs P18–20 (p<0.0001, N=23 P11–14, N=24 P18–20) comparisons (Dunn’s multiple comparisons test). Bars represent mean ± S.E.M. and individual data points are the total switches for 20 mins for each rat pup group.

To account for the probability of staying in a given state, we then analysed data per second of the experiment while taking into account transition stays (for example, staying in aggregon 0 for 5 seconds would make P_0-0_ probability of 1). We then created a three-state Markov chain model which accounted for the probability of state-stays (**Figure S1A**). We then took the mean for each state across groups at each developmental state to compile a probability matrix. Across all developmental stages, we see that the probabilities of staying in a given aggregon were the highest per second events (diagonal of matrix) and the probabilities of transitions between states were lower (**Figure S1B**). To compare the model to the data, we ran the Markov model 25–30 times from a starting value of 1 (aggregon 0) with 1200 steps to simulate the observations made of the huddling test. In all age-groups we observed a rise of the aggregon state across time, which stabilised in last portion of the test, both for the data (colored lines) and the model (black) (**Figure S1C, left**). The rise time of the aggregon score appeared faster in the model, however the final aggregon score reached was not different between the model and data (**Figure S1C, right**). Thus, we find that huddling behaviour can be predicted with state probabilities that are changed in development. Staying in each state is more common per second than transitioning; transitions between states (switches) increase in development.

### Aggregon durations and switches are not influenced by kinship

We hypothesized that huddling dynamics would be influenced by kinship, thus we analysed data while separating kin and non-kin group data. We found no kinship differences in aggregon durations across developmental ages (**Figure 4A**). We additionally analysed the number of aggregon switches between kin and non-kin groups and found no statistical differences (**Figure 4B**). Thus, huddling dynamics are strong and consistent in kin and non-kin groups, without kinship differences, thus rejecting our hypothesis. We additionally examined whether huddle dynamics differed in male and female animals and found no sex-differences in huddling behaviour. Thus, the drive to huddle seems to override preferences based on sex and kinship.

**Figure 4.**
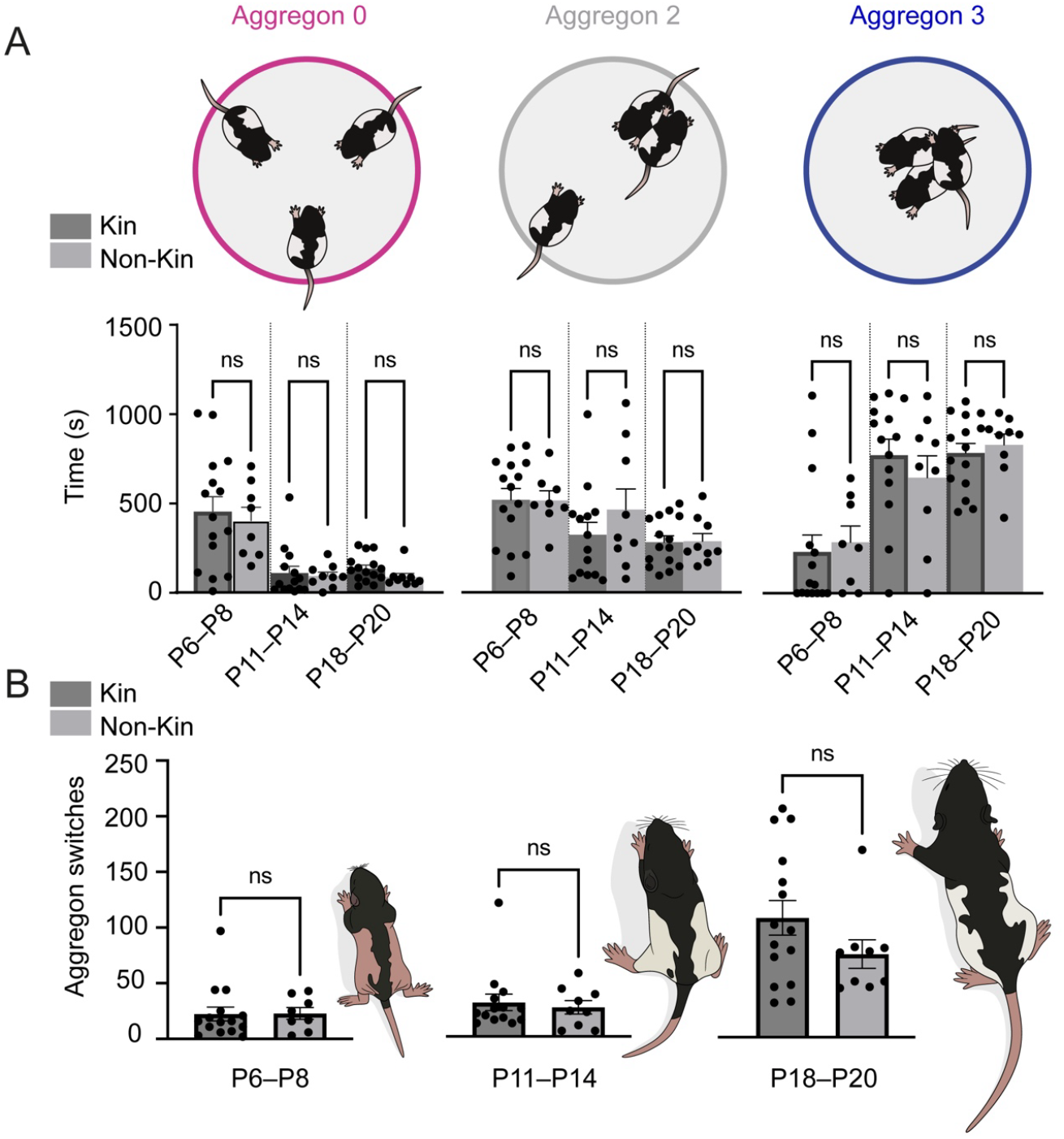
Huddle aggregon times and switches are not different for kin and non-kin huddle groups. A) Time spent in aggregons did not differ between kin and non-kin huddle groups. Aggregon 0 (P6–8, N kin = 15, N non-kin = 8 p = 0.7763; P11–14, N kin = 14, N non-kin = 9, p = 0.557; P18–20, N kin = 15, N non-kin = 9, p = 0.096), aggregon 2 (P6–8, N kin = 15, N non-kin = 8, p = 0.825; P11–14, N kin = 14, N non-kin = 9, p = 0.403; P18–20, N kin = 15, N non-kin = 9, p = 0.907), aggregon 3 (P6–8, N kin = 15, N non-kin = 9, p = 0.374; P11– 14, N kin = 14, N non-kin = 9, p = 0.344; P18–20, N kin = 15, N non-kin = 9, p = 0.770). B) Number of aggregon switches did not differ between kin and non-kin huddle groups P6–8 (N kin = 15, N non-kin = 8, p = 0.602), P11–14 (N kin = 14, N non-kin = 9, p = 0.865), P18–20 (N kin = 15, N non-kin = 9, p = 0.221). Comparisons made with Mann-Whitney test. Bars represent mean ± S.E.M..

### Aggregon complexity is inversely related to the amount of calls

To examine the interaction between aggregons and ultrasonic vocalisations (USVs), we synchronised vocalisations to align to the video recording of huddling. The timing of each vocalisation was detected with DeepSqueak and manually curated. When rat pups formed aggregons of increasing size, the number of vocalisations reduced, in all age-groups (**Figure 5A**). We observed that aggregon score per minute and the number of calls per minute followed a linear trend where calls reduced as aggregon number increased, with the most negative slope found in P6 – 8 and P11 – 14 groups (**Figure 5B**). Across development, the spectral characteristics of calls changed from being uniform in principal frequency to more complex with age, particularly in P18 – 20 groups (**Figure S2A**). The peak frequency of the vocalisation spectrogram was significantly higher when comparing P11 – 14 and P18–20 groups, likely reflecting the emergence of higher frequency calls in P18–20 animals (**Figure S2B**). The effect of vocal quieting with huddling was most pronounced in youngest age groups (Figure 5B: P6– 8 R^2^ = 0.93, slope = -98.28; P11–14 R^2^ = 0.91, slope = -103.4; P18–20 R^2^ = 0.58, slope - 17.09), thus the quieting effects may be most pronounced in lower peak frequency calls such as those elicited in separation and distress conditions.

**Figure 5.**
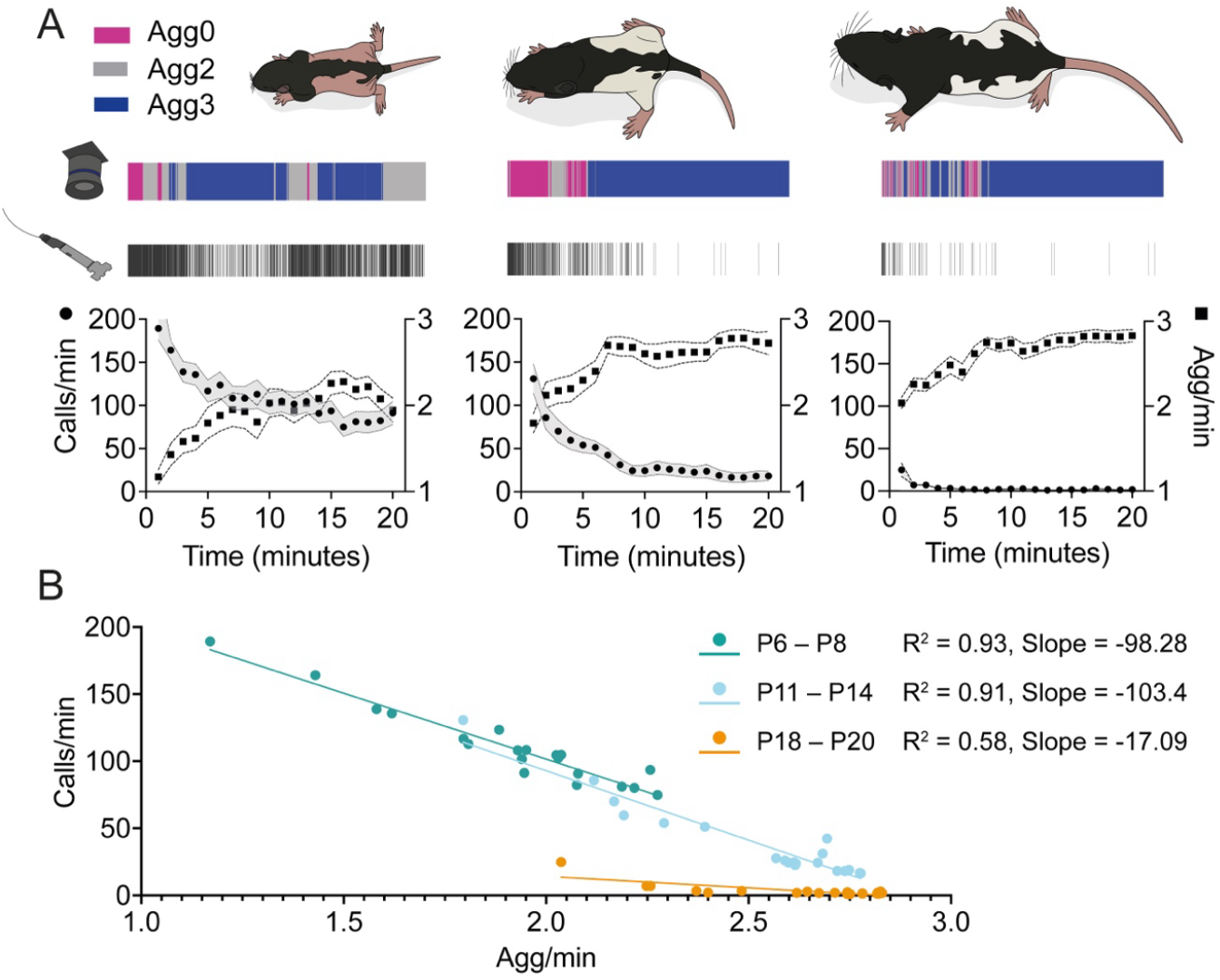
Vocal quieting occurs throughout the duration of huddle observation and is inversely related to aggregon score, in all pre-weaning age-groups. A) Examples of aggregon events aligned to vocalisations at P6 – 8 (top), P11 – 14 (middle) and P18 – 20 (bottom). Magenta bars indicate aggregon 0, gray indicates aggregon 2 and blue aggregon of 3. Ultrasonic vocalisation (USV) rate calculated per minute for each group at P6 – 8 (left), P11 – 14 (middle) and P18 – 20 (right). Squares indicate the mean aggregon score per minute across all groups, dotted line on either side indicates the mean ± SEM. Circles indicate the mean number of calls for each minute across all groups, shaded area indicates the mean ± SEM. B) Mean call rate for each minute vs mean aggregon score per minute (P6 – 8 N=18, R^2^ = 0.93, slope = -98.28; P11 – 14 N=19 R^2^ = 0.91, slope = -103.4; P18 – 20 N = 22, R^2^ = 0.58, slope -17.09). Group data only included if both video and USV recordings collected with TTL alignment signal intact (two groups lacked alignment signal P18 – 20).

### Ultrasonic vocalisations are reduced in young (P6 – 8), but not older kin groups

To address whether kinship differences could be observed in the huddle task, we examined the mean number of calls per minute for kin and non-kin groups. We observed that the mean number of calls per minute was reduced in youngest kin groups (P6–8). The number of calls per minute did not differ in kin vs non-kin groups in older (P11–14 and P18–20) groups (**Figure 6**).

**Figure 6.**
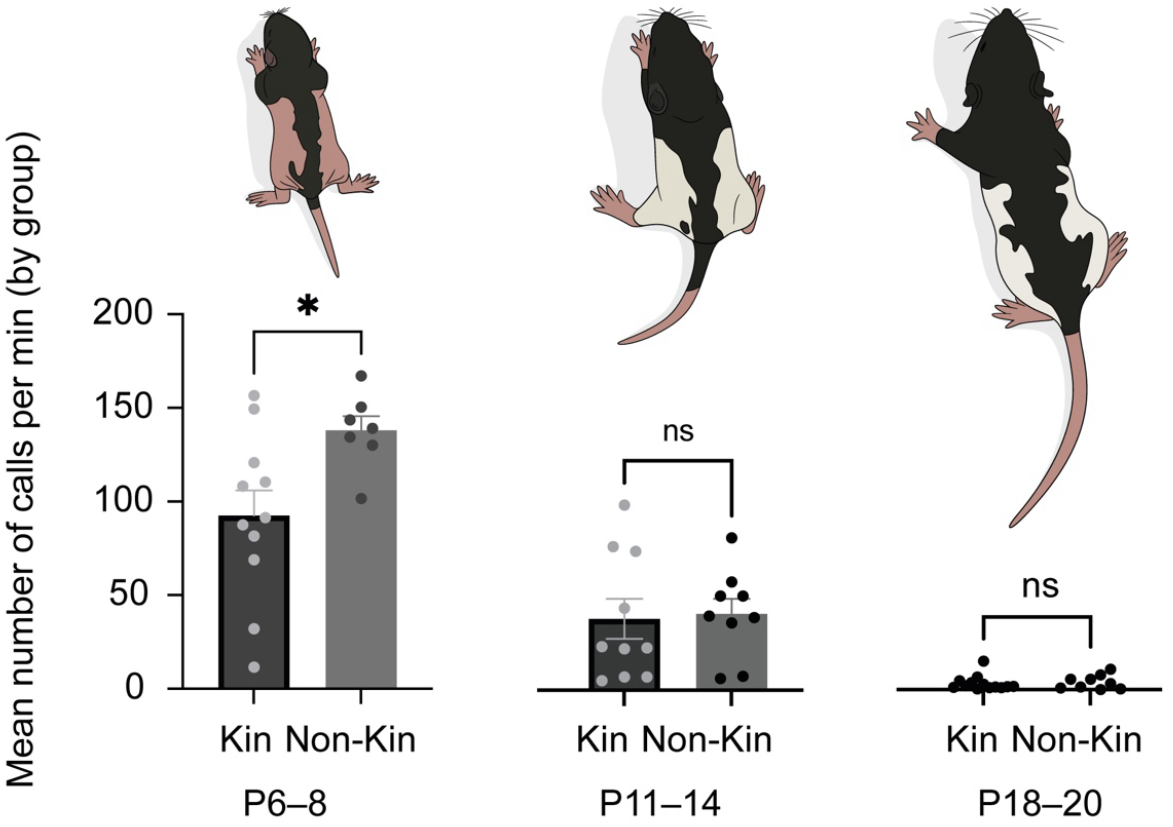
Ultrasonic vocalisations (USV) in huddle trial are reduced in young age (P6– 8) kin vs non-kin groups, but not older pre-weaning age groups. The average amount of calls over the 20 minute recording was calculated for each group across developmental age. Left: Kin (N = 11 groups) vs Non-kin (N = 7 groups), p = 0.0268. Middle: Kin (N = 10 groups) vs Non-kin (N = 9 groups), p = 0.6038. Right: Kin (N = 13 groups) vs Non-kin (N = 9 groups), p = 0.948. Mann-Whitney test. Left axis corresponds to all graphs. Bars indicate mean ± SEM.

### Huddle touch exclusion increases vocalisations in early-mid pre-weaning kin age groups

We lastly examined the role of touch in vocal quieting and kin-dependent huddle vocalisations. Thus, we designed a divider which separated touch between the three pups in the huddle trial. Airflow was not contained; thus pups were expected to maintain abilities to smell each other. The huddle trials were carried out with the same protocol as in previous experiments (**Figure 7A**). When sibling (kin) groups without touch (divider intact) were compared with kin groups without divider, we observed increased vocalisations without sibling huddling touch in P6–8 and P11–14 groups, but not in P18–20 groups (**Figure 7B**). Thus, social touch appears to be an important regulator of intrinsic state in early pre-weaning huddling behaviour.

**Figure 7.**
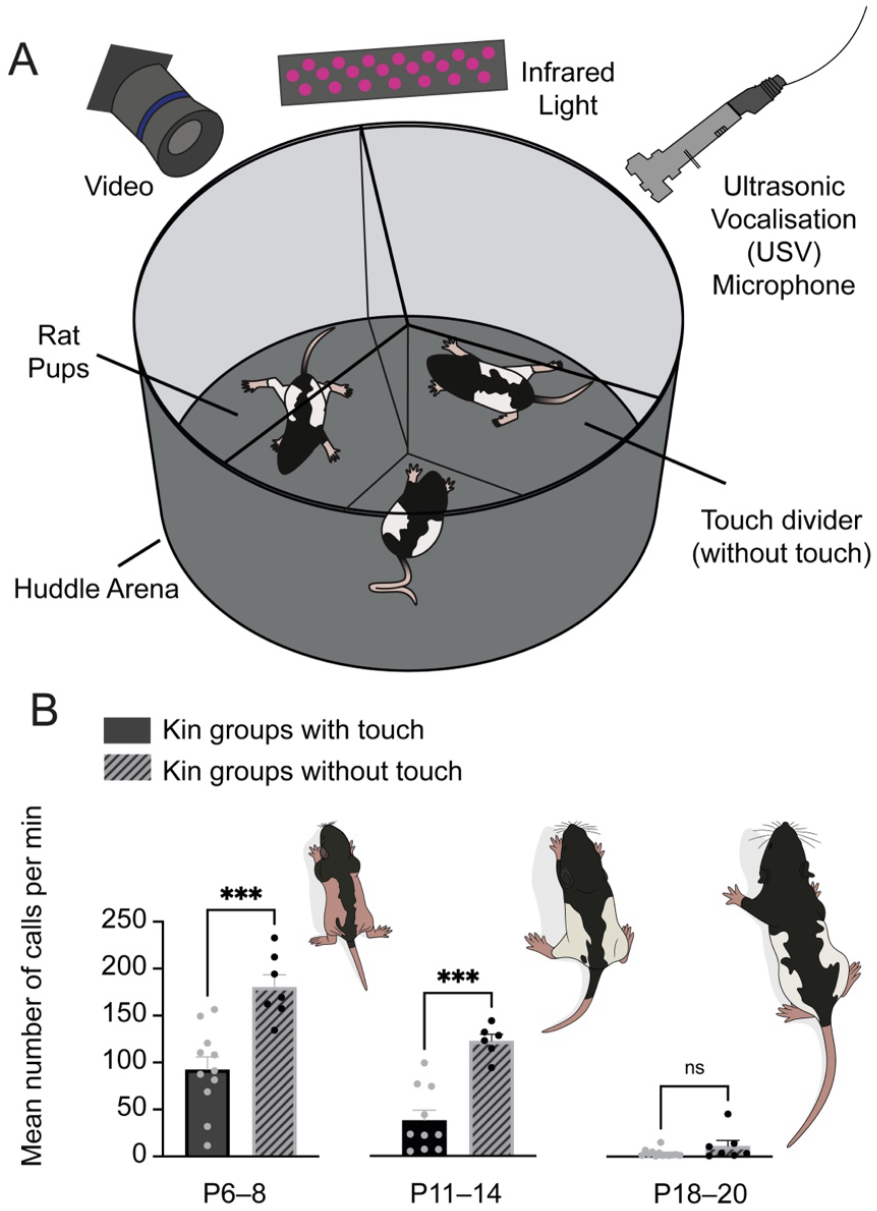
Huddle touch exclusion increases vocalisations in early-mid pre-weaning development. A) Sibling rat pups were placed in the huddling arena with a 3- chamber divider separating huddling touch. The experiment was conducted for 20 minutes, as in previous huddling tests. B) The average amount of calls per minute, per group was higher in kin groups with the touch divider present (no-touch) compared with kin groups which were free to touch. P6 **–** 8 (p = 0.0003, N kin touch = 11, N kin no-touch = 7), and P11 **–** 14 (p = 0.0005, N kin touch = 10, N non-kin touch = 6) but not at P18**–**20 (p = 0.948, N kin touch = 13, N non-kin touch = 9). Bars indicate mean ± SEM.

## Discussion

We report that huddling behaviour 1- shows age-dependent changes in the dynamics of huddle formation, 2- huddle aggregon dynamics are not dependent on kinship 3- aggregon complexity is correlated with vocal quieting 4- USV reduction in kin-groups at the youngest age suggest a more calm internal state in the huddle task, 5- absence of touch in the huddle task increases USVs in P6–8 and P11**–**14 groups.

### Huddle aggregon complexity and dynamics with development

We observed that rat pups spontaneously form huddles without training. In the earliest stage of development (P6–8), the time spent in lower aggregon states may be due to under- developed motor abilities. Thus, rat pups of this age rely more on the parent to move them together in the nest. Interestingly, although thermo-regulatory need for huddling decreases in development [11], rat pups continue to form huddles with increasing speed (**Figure 2**). Furthermore, we continue to observe vocal quieting in older pups, indicating that the motivation to huddle may be a long-lasting mechanism for social bonding which is rewarding and calming.

### Huddle drive overrides kinship selectivity

Despite previous reports of kinship differences in huddling behaviour [12], our measures of huddling aggregon duration and switching did not reveal kinship differences (**Figure 4**). Similarly, although indications in the literature pointed towards sex-differences in huddling, we did not observe this. One possibility for this is the design of the huddling test. In Hepper’s work, groups of kin and non-kin rat pups were mixed and scored based on fur-markings, thus kin differences in huddling may arise when presented with the simultaneous choice of huddling with kin or non-kin. We chose a simpler design, where we could be certain of the composition of the rats being studied. Furthermore, with mixed kin and non-kin groups, the assessment of vocal behaviour would not be distinguishable between kin and non-kin. Further studies could implement Hepper’s approach of mixing groups, wherein the pups would be given the choice of kin and non-kin to associate with in the huddle group. Alternatively, it may be that the instrinsic drive to huddle overrides kinship selectivity. The rewarding aspects of sharing of warmth and touch for rat pups may be sufficiently strong that the behavioural dynamics are indistinguishable for related and unrelated conspecifics.

### Vocal quieting with aggregon formation

We observed consistent reduction in the number of vocalisations as rat pups huddled more (**Figure 5**). Vocal quieting was consistent although the spectral characteristics of the calls showed developmental changes (**Figure S2**). Thus, regardless of the vocalisation type, the overall tone of vocalising- or not vocalising reduces and is associated with increased contact. Contact quieting has been observed in other social behavioural paradigms [14,19] and may be the reason for what we observe. Many studies report contact quieting between the pup and caregiver [14,18,19], our study confirms that siblings provide potent calming and quieting social stimuli as well.

### Young-pup kin vocal quieting and dependence on touch

We found that despite indistinguishable huddling dynamics in kin and non-kin pup groups, calls were reduced in sibling huddle groups in the youngest developmental age (P6–8), but not in older pup groups. Thus kinship, familiarity, or a combination of the two factors may calm sibling pups in the huddle task. Indeed, it is known that anxiolytics reduce USVs in infant rats [20]. Although the recognition of kin is likely to be regulated by olfactory stimuli [21–23], when we removed touch in the huddling task, the amount of calls was no longer reduced. The touch- dependence of vocal quieting was observed in the P6**–**8 and P11**–**14, but not in older (P18**–** 20) pups (**Figure 7**). Thus, although recognition of siblings may be calming, the intact ability to touch and huddle is necessary for this potentially anxiolytic stimulus to be maintained.

### Neural mechanisms and social homeostasis

Recent work has elegantly shown that vocal quieting with social reunion occurs in adult mice and the overall level of ongoing vocalisation is heightened in groups which have undergone social isolation. The group further shows that tactile stimulation is key to social rebound and that neurons of the medial pre-optic nucleus (MPN) of the hypothalamus underly tactile social homeostasis [24]. Another recent analysis of huddling in mice, suggests a role for the dorsomedial prefrontal cortex (dmPFC) in active and passive huddling decisions [25]. Work in Prairie voles implicates corticostrial circuits in affiliative partner bonding [26]. Future studies in a variety of rodent species will address if the underlying brain mechanisms are conserved.

Indeed, work from our group on the development of kin preference in rat pups point to the lateral septum (LS) as a target area for examination in huddling behaviour [21,27]. Other recent work in mice demonstrates that neurons of cortex which project to the LS underly the development of social novelty preference [27]. Work in Spiny mice similarly implicate cortical to LS circuitry as a mediator of group size preference [28]. Indeed, the LS may constitute an evolutionarily conserved hub which orchestrates social group preference and the underlying emotional states.

## Conclusions

Overall, we have shown that huddling behaviour is a natural and self-organising behaviour in rat pups. Unique dynamics of contact are observed with development and vocal quieting with aggregon complexity occurs in all age-groups. Intrinsic states, indicated by USV analysis, indicate that kinship and sibling touch are calming in the youngest developmental time periods. We suggest that huddling is a calming and rewarding activity amongst siblings and un-related conspecifics which balances social homeostasis in development, through an interplay of tactile, vocal and kinship dynamics.

## Acknowledgements

We thank Lynn Morrison, Neil Odey and Callum Davidson for technical support. Thanks to Connie Yung, Grace Chattey, Zuzanna Baran and Isabel Ibeson for helpful discussion. Gratitude to Víctor Angulo for rat drawings. This work was supported by the Simons Initiative for the Developing Brain (SIDB), the University of Edinburgh MSc programme in Integrative Neuroscience (HT) and a Simons ESAT fellowship (AC).

---

**Figure S1.**
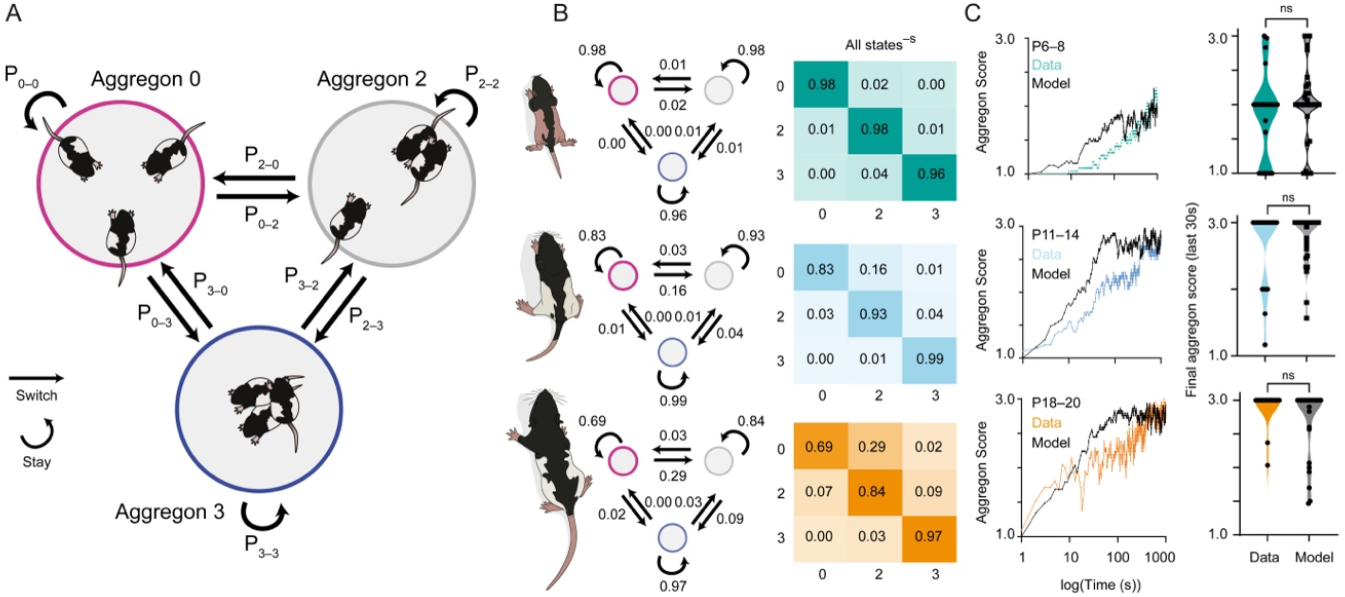
The dynamics of huddle state transitions are regulated by developmental age. A) The probability of huddle aggregons states and transitions between aggregon states were calculated. B) The probability of each state transition was calculated for the groups in which aggregon data and vocalisations could be aligned (P6 – 8 N = 22, P11 – 14 N = 20, P18 – 20 N = 21). Throughout the 20-minute experiment, 1200 states were given for each rat pup group, the probability of each transition was then calculated per group and a mean was taken across the groups. Left: For each age-group the probabilities for each state. Right: Probability matrix, where each row sums to 1. C) Left: The model was run 25 – 30 times across 1200 steps, simulating the length of the huddling experiment. The data of the aggregon scores (coloured lines) are plotted on the model output (black). Right: the mean aggregon score was calculated for the last 30 seconds of the data and the model. The final aggregon score did not differ between the data and the model for any age-group (P6 – 8, p = 0.134; P11 – 14, p = 0.894; P18 – 20, p = 0.085; Mann Whitney test).

**Figure S2.**
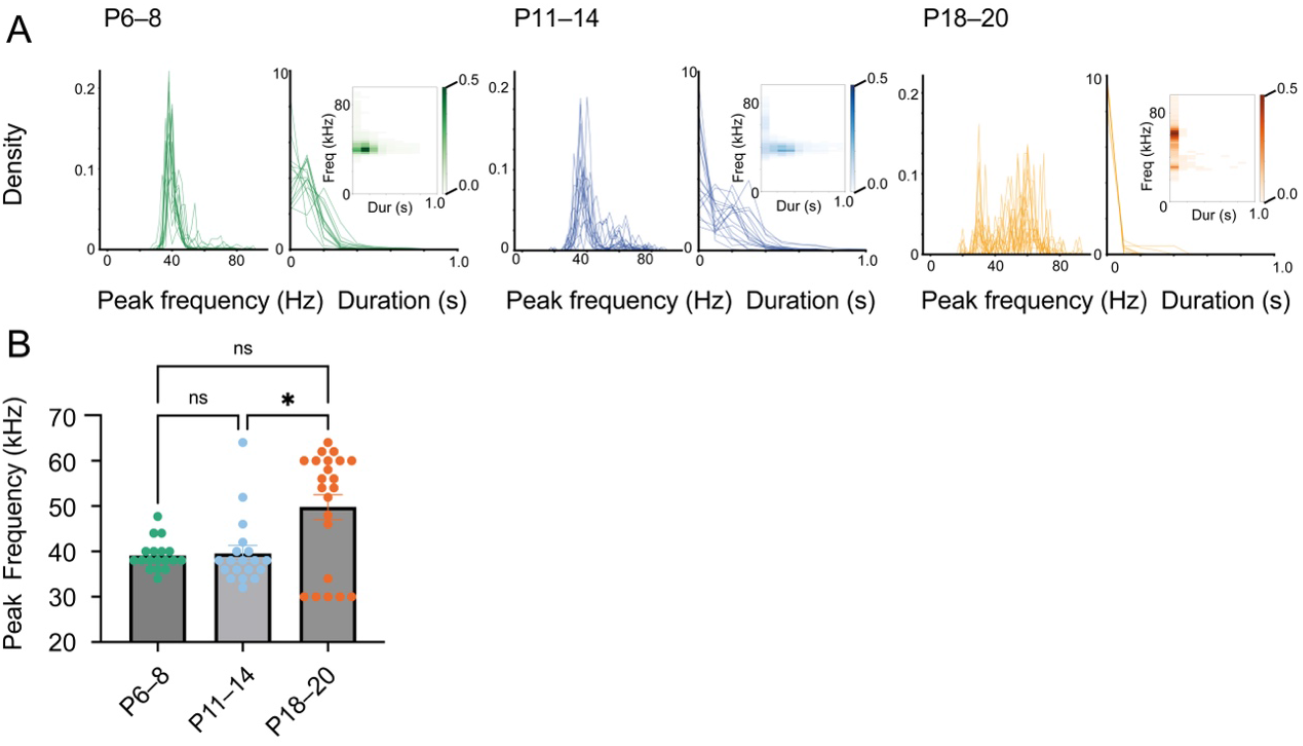
Vocalisation spectral features change in development. A) (left) probability density of the peak frequency (Hz) and duration (s) of calls at P6 – 8, (middle) at P11 – 14 and (right) at P18–20. All calls included in 20 minute huddle trials. B) Comparison of mean peak frequency for each groups of rats across development. Peak frequencies were found to vary across age (Kuskall-Wallis test p = 0.0276, N P6 – 8 = 18; N P11 – 14 = 19 ; N P18 – 20 = 22) with multiple comparisons revealing no significant difference P6 – 8 vs. P11 – 14 (p > 0.999) or P6 – 8 vs. P18 – 20 (p = 0.1246). A significant difference was found P11 – 14 vs. P18 – 20 (p = 0.0397). Bars indicate mean ± SEM.

